# Heat stress and the temporal dynamics of insect growth

**DOI:** 10.1101/2023.09.06.553041

**Authors:** Joel G. Kingsolver, J. Gwen Shlichta, M. Elizabeth Moore

## Abstract

Temperature affects rates of ingestion, growth and other key processes in insects and other ectotherms, a relationship described by the thermal performance curve. But the history of temperature exposure can cause stress or acclimation responses, altering the relationship between current temperature and performance. The temporal dynamics of such time-dependent effects for ectotherm growth are poorly understood. We quantify how growth and ingestion change over time at different constant temperatures during the final larval instar for two thermal-generalist insect species, *Pieris rapae L.* and *Manduca sexta L*. Initial growth rates were greatest at higher temperatures for both *P. rapae* (29-35 °C) and *M. sexta* (34-38 °C); but growth rates at higher temperatures declined over time. As a result, the optimal and maximal temperatures for growth declined over time (2-4 days) in both species. Unlike *P. rapae*, *M. sexta* could maintain growth and survival to wandering and pupation at temperatures above 35 °C. For *M. sexta* larvae at 40 °C, ingestion continues even as growth declines, suggesting that post-ingestive processes limit growth rates at these high temperatures. Our results show how time-dependent effects at higher temperatures develop over hourly to daily time scales, with important consequences for understanding and modeling organismal responses to heat waves and extreme temperature events.

## Introduction

The growth and developmental rates of insects and other ectotherms depend strongly on their body temperatures. The thermal sensitivity of an organism is often characterized in terms of a thermal performance curve that relates organismal performance to body temperature (Huey and Stevenson 1979, Huey and Kingsolver 1989). Thermal performance curves for rates of growth, development and most other biological processes are non-linear, with an intermediate optimal temperature for performance, and reduced performance at both lower and higher temperatures (Dell et al. 2011). The effects of the non-linearity of performance curves for mean performance in variable environments have been thoroughly explored theoretically and empirically (Worner 1992, Lactin et al. 1995, Ruel and Ayres 1999, Tobin et al. 2001, Colinet et al. 2015); and the shapes of performance curves have important consequences for predicting responses to climate change (Deutsch et al. 2008, Clusella-Trullas et al. 2011, Kingsolver et al. 2013, Sinclair et al. 2016).

The use of performance curves assumes that performance depends only on current environmental conditions. However, many numerous studies show that the history and pattern of prior thermal exposure can influence performance (Schulte et al. 2011, Mislan et al. 2014, Rezende et al. 2014). These effects can be particularly important at higher temperatures. For example in insects, increasing duration of exposure to temperatures in the upper thermal range can reduce performance, even at non-lethal temperatures As a result, using thermal performance curves based on data from constant-temperature experiments can produce poor predictions about mean performance in diurnally fluctuating conditions (Niehaus et al. 2012, Kingsolver et al. 2015, Sinclair et al. 2016).

The effects of duration and pattern of thermal exposure on performance—called ‘time-dependent effects’ (Kingsolver et al. 2015)-- can result from stress and acclimation (Hoffmann and Merila 1999, Sorensen et al. 2003). Stress and acclimation responses to temperature have been widely documented in many organisms, but the dynamics of how performance changes over time at different temperatures are not well-known. Quantifying these temporal dynamics is important for understanding how stress and acclimation can change the shape and position of thermal performance curves (Kingsolver and Woods 2016).

In this study, we quantify how temperature affects the temporal dynamics of growth and ingestion during the final larval instar of two insect species, *Pieris rapae L.* and *Manduca sexta L*. These widespread agricultural pests can grow and develop over a wide range of temperatures, but they differ in their performance and tolerance at high temperatures. By rearing larvae in fluctuating temperatures and then exposing them to different, constant temperatures throughout the final instar, we can evaluate how exposure duration affects rates of growth and ingestion at different temperatures, and how this causes temporal changes in thermal performance curves. By examining ingestion in addition to growth during these exposure periods we can directly compare how growth and ingestion rates changed with time across different temperatures, and how performance curves for ingestion rate change with time.

## Materials and Methods

### A. Study systems

#### 1. Pieris rapae

The Imported Cabbageworm *P. rapae* was originally native to Europe but is now found throughout most of North America and other continents (Scudder 1887). Its larval host plants include a variety of wild and domesticated forms of *Brassica* and related genera in the family Brassicaeae; in many regions it is agricultural pest on cabbage, kale, collards and other domesticated forms of *Brassica oleracea* L. *Pieris rapae* has five larval instars, and more than 70% of total larval growth (mass increase) occurs during the final instar (Kingsolver et al. 2006). Previous studies suggest that the effects of temperature on growth rate are similar for different instars (Chen and Su 1982, Kingsolver 2000). Our lab and field observations indicate that *P. rapae* larvae do not actively regulate body temperatures except to avoid extreme high temperatures, and they will feed and grow in both light and dark periods under appropriate thermal conditions (Kingsolver et al. 2004).

Short-term (4-24h) larval growth rates are fastest at body temperatures near 35 °C (Kingsolver et al. 2004, Izem and Kingsolver 2005), but larvae experience increasing mortality rates at chronic constant temperatures above 30-32 °C, with complete mortality at 35 °C (Kingsolver 2000). Field measurements in collard (*B. oleracea*) gardens in Seattle, Washington (USA) show that the body temperatures of individual larvae may vary by more than 20 °C during a single diurnal cycle, and that an individual larva may experience body temperatures spanning the full temperature range allowing effective growth (15- 35 °C) during its larval life (Kingsolver 2000, Kingsolver et al. 2004). Our studies here were initiated with female *P. rapae* collected from organic farms near Seattle WA in Sep 2001.

#### 2. Manduca sexta

The Tobacco Hornworm *M. sexta* is native to the Americas, and occurs across Central America, northern South America, and southern North America. Larvae feed primarily on host plants in the family Solanaceae, including domesticated crops such as tobacco and tomato. *Manduca sexta* typically have five larval instars and more than 80% of mass increase occurs during the final instar. Towards the end of the final instar, larvae stop feeding and wander off the host plant to pupate in the soil. Mass at wandering is strongly correlated with pupal mass, adult size and female fecundity (Davidowitz and Nijhout 2004, Diamond and Kingsolver 2010a, b). The experiments described here were conducted in spring and summer 2016, and used *M. sexta* larvae reared on a standard artificial diet (Bell and Joachim 1976) from a laboratory colony at UNC maintained at 25-26 °C. This colony was originally established from field populations near Raleigh NC in the 1960s.

Short-term (4-24h) growth rates of *M. sexta* larvae are fastest at body temperatures of 34-36 °C, but under chronic, constant temperatures larval growth rates and survival decline rapidly above 30 °C (Kingsolver and Woods 1997, Kingsolver and Nagle 2007, Kingsolver et al. 2015). Field studies in both the southeastern and southwestern US show that larvae regularly experience body temperatures exceeding 35 °C during the summer months (Casey 1976, Diamond and Kingsolver 2010a, Kingsolver et al. 2012). Laboratory colony and field populations of *M. sexta* differ in maximal larval growth rates and heat tolerance, but they have similar growth responses to alternating rearing temperatures (Kingsolver and Nagle 2007, Kingsolver et al. 2009, Diamond and Kingsolver 2010b).

### B. Experiments

#### 1. Growth in *P. rapae*

Studies with *P. rapae* began with adult females collected from organic farms near Seattle WA. Each female was allowed to lay eggs on collard plants (*Brassica oleracea* var. collard, Champion variety) in an individual oviposition cage in the greenhouse. After hatching, all larvae were fed fresh collard leaves in petri dishes in an environmental chamber (Percival 36VL), under a diurnally fluctuating (11-35 °C) thermal regime and 16hL:8hD photocycle that mimic typical field conditions for July-August in Seattle (Kingsolver 2000).

The experiment began with newly molted 5th instar larvae. Sets of 15-30 5^th^ instar larvae, representing the offspring from several different females, were randomly assigned to one of six constant test temperatures in environmental chambers: 11 °C, 17 °C, 23 °C, 29 °C, 35 °C, or 40-41 °C. The initial body mass of the caterpillar was measured, and the larva was kept in an individual petri dish with fresh, young collard leaves and a damp sponge to reduce water loss from the leaves. The mass of each larva was then measured to ± 0.01mg (Mettler Toledo AT-261 balance) at 5 times: 6h, 24h, 30h, 54h, and 80h. At the lowest temperature (11 °C), each larva was measured at 48h rather than 30h or 54h; at the highest temperature (41°C), each larva was measured at 8h rather than 6h (see below). Fresh collard leaves were added at each time. We also noted whether active feeding (visible leaf consumption) occurred during each time interval, as well as any mortality. The results reported here are for larvae that survived and fed actively during the time period of interest.

#### 2. Growth and ingestion in *M. sexta*

Studies with *M. sexta* began with eggs obtained from our laboratory colony at UNC. Our basic protocol follows that used by Kingsolver et al. (2015). Larvae were reared from hatching on artificial diet in a programmable environmental chamber (Percival 36VL) with a long-day (14hL:10hD) photocycle and a diurnally fluctuating thermal regime (11-35 °C). Larvae were reared at low density in bins containing abundant diet that was changed every two days.

On the morning following molt into the 5^th^ (final) larval instar, each larva was placed in an individual petri dish with a block of diet; the initial mass of each larva and of its block of diet was measured. The size of the diet block differed among test temperatures so that sufficient food was available for the duration of the measurement time period. Sets of 15-30 larvae were randomly assigned to one of five test temperatures in different environmental chambers: 25 °C, 30 °C, 34 °C, 38 °C or 40 °C. We did not consider lower test temperatures because previous studies did not show time-dependent effects at lower temperatures (Kingsolver et al. 2015). The mass of each larva was measured at 8 times: 6h, 24h, 30h, 48h, 54h, 72h, 96h, 120h and 144h. At each time point the remaining diet block in the dish was weighed, and a fresh diet block was weighed and added to the dish. During each time interval we also noted whether active feeding (visible consumption) occurred, as well as any mortality. Larvae that had begun to wander (defined by the appearance of the dorsal vessel) were weighed, removed and placed in wooden pupation chambers; survival, date and mass at pupation were also recorded. The results reported here are for larvae that survived and fed actively during the time period of interest.

Because mass change of each diet block may reflect both larval ingestion and water loss from the block, we quantify ingestion in terms of dry mass of each block. We placed fresh diet blocks in each of the test temperatures, and measured fresh and dry (after 3 days in a drying oven at 58 °C) mass of blocks for time intervals of 0h, 6h, 18h and 24h (ten blocks for each time interval for each test temperature). We used these data to estimate % water content of diet at each temperature at each time interval, and to quantify ingestion as the change in dry mass of a block over the time interval.

### C. Analyses

One main goal of our experiments was to quantify growth curves-- changes in body mass over time—during the final larval instar, and how these vary across different constant test temperatures. Because growth during the 5^th^ instar is approximately isometric (rather than allometric or exponential) for *M. sexta* (Davidowitz et al. 2005, Nijhout et al. 2006, Sears et al. 2012) and *P. rapae* (see Results), we modeled mass on an arithmetic (rather than log) scale. In the figures (Figs. 1 and 3) we present data on larval mass over time until the cessation of feeding (*P. rapae*), wandering (*M. sexta*), or death; but we restrict statistical analyses to the time period of active growth (see Figs. 2 and 4). Hence we omitted from the analysis the mass measurement for any larva that had stopped feeding or started wandering. Note that larvae with relatively slow growth and development rates will complete their growth at later times than rapidly developing larvae, biasing the mean growth curve at later times (see Figs. 1-2 and 3-4); to reduce this bias we restricted the time period in our analyses to 0-54h for *P. rapae* and 0-144h for *M. sexta* (see Results and Discussion).

**Figure 1.**
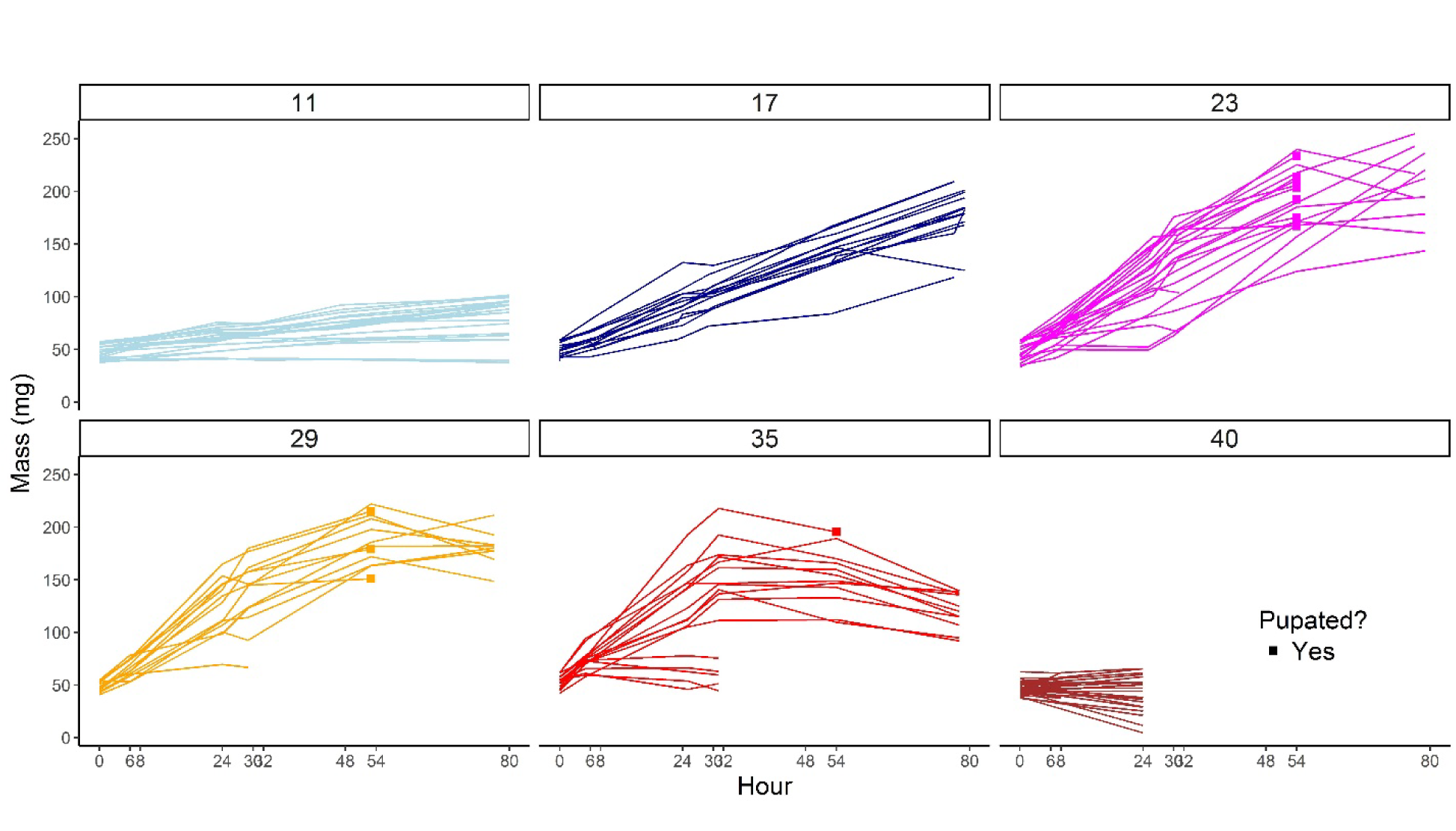
Individual growth curves throughout the 5^th^ instar of *Pieris rapae* by test temperature. Mass at time 0h represents the initial mass upon molting into the final 5^th^ instar. Individuals who successfully made it through the 5^th^ instar to pupation during the time that measurements took place are denoted with a square.

**Figure 2.**
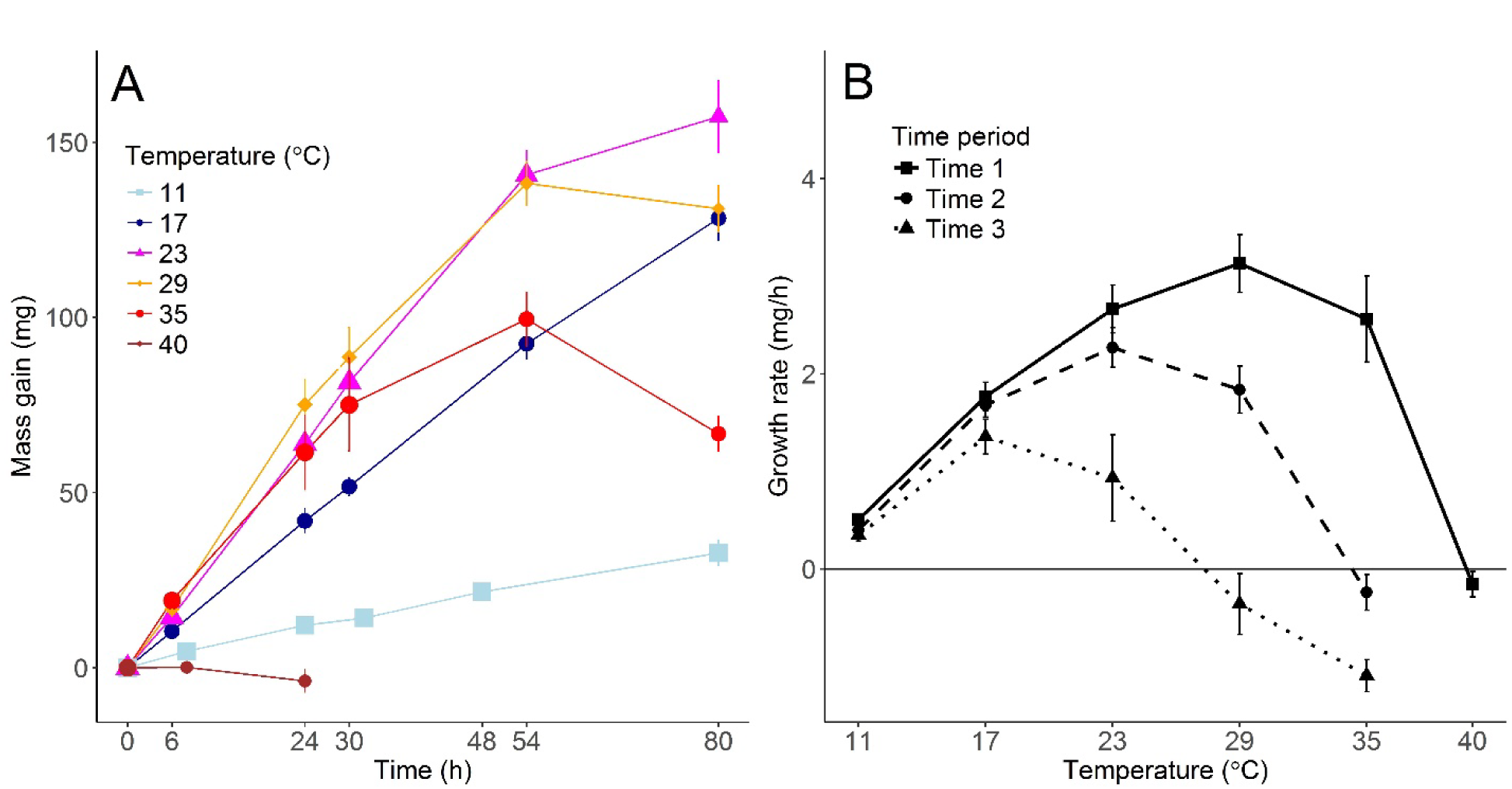
Mean mass gain (mg) with standard error during the 5^th^ instar by time (A) and mean growth rate (mg/h) with standard error by test temperature (B) for *P. rapae*. Line types denote 3 different time periods of sampling. Period 1 represents Time 0-24h of the 5^th^ instar for all test temperatures, Period 2 represents Time 24-48h or Time 30-54h depending on the test temperature, and Period 3 represents Time 48-80h or Time 54-80h depending on the test temperature. All measurements prior to pupation are included in the mean calculation.

**Figure 3.**
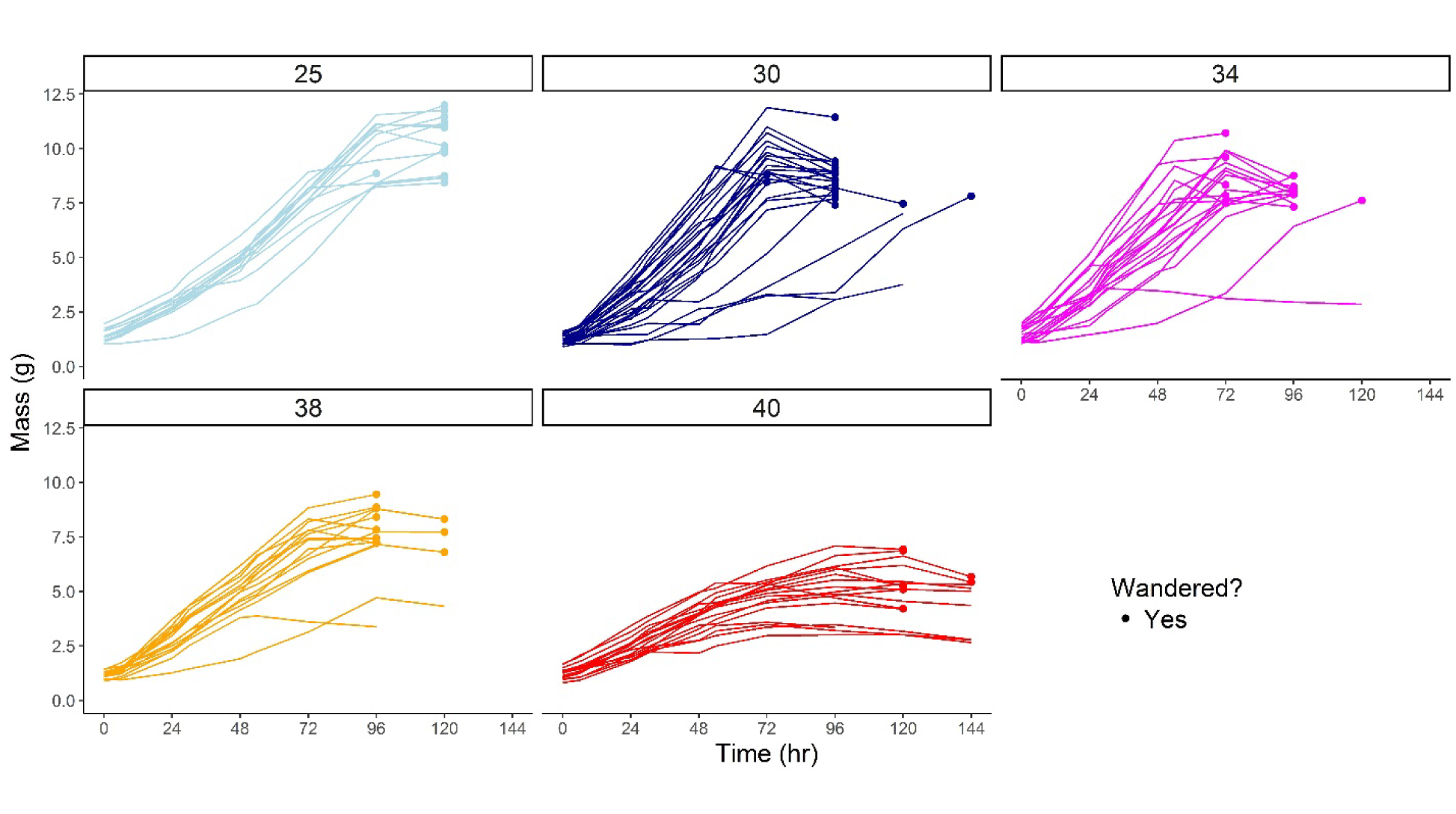
Individual growth curves throughout the 5^th^ instar of *M. sexta* by test temperature. Mass as time 0h represents the initial mass upon molting into the final 5^th^ instar. Individuals who successfully made it through the 5^th^ instar to the wandering stage during the time that measurements took place are denoted with a circle.

**Figure 4.**
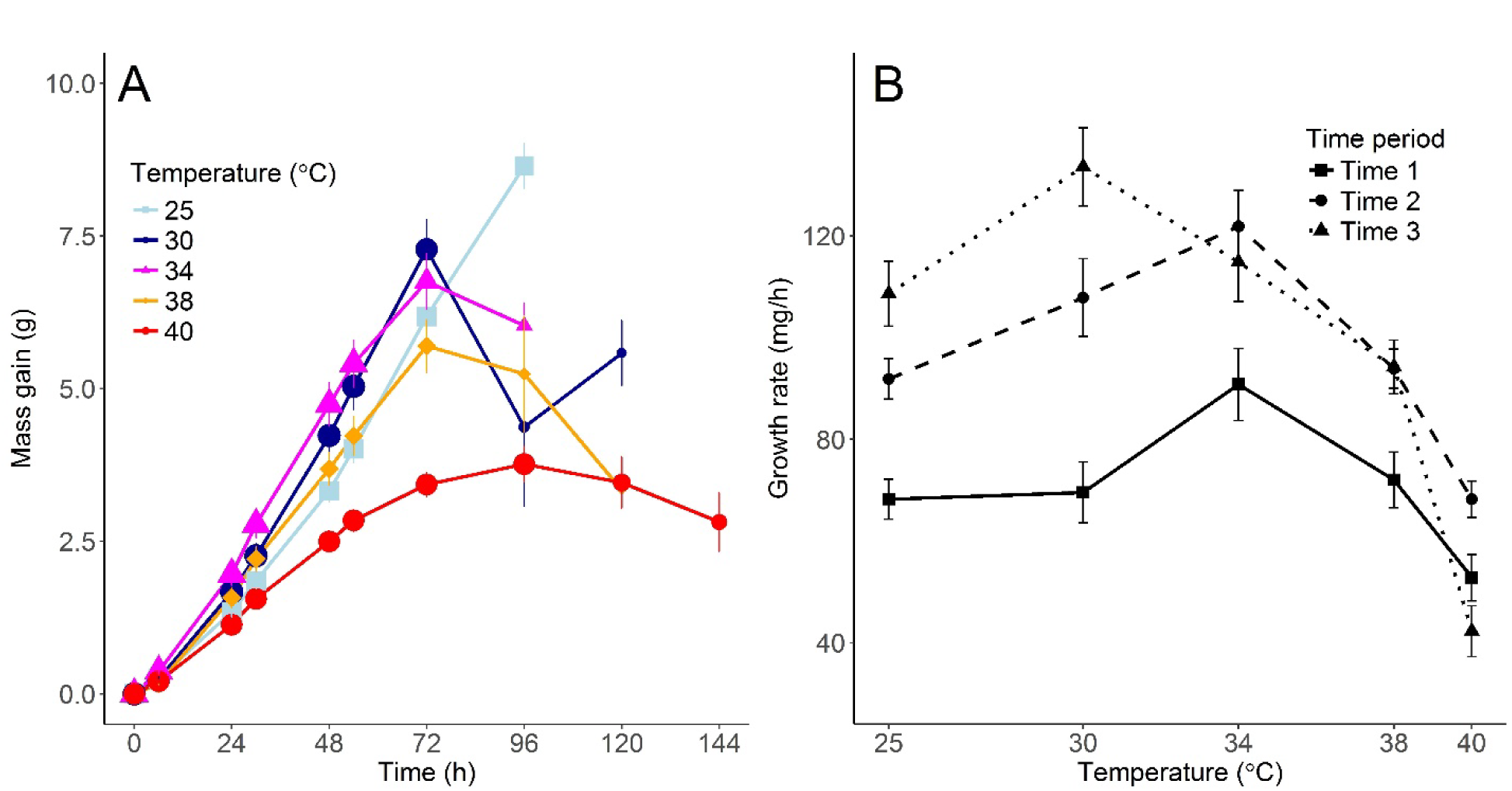
Mean mass gain (g) with standard error for during the 5^th^ instar by time (A) and mean growth rate (mg/h) with standard error by test temperature (B) for *M. sexta*. Line types denote 3 different time periods of sampling, including Time 0-24h, 24-48h, or 48-72h of the 5^th^ instar. All measurements prior to wandering are included in the calculation of the means.

We used linear mixed-effects models (function lme in R library nlme) to analyze larval mass (m). We considered time (t) and test temperature (T) as fixed effects; time was modeled as a 2^nd^ order (*P. rapae*) or 3^nd^ order (*M. sexta*) polynomial, and temperature was included as a factor. Individual was included as a random (intercept) effect in the model. We used maximum likelihood (rather than REML) to fit the model, because we are primarily interested in the fixed effects. Note that the rate of change in mass (dm/dt) is the growth rate, so that negative curvature (the 2^nd^ order term, t^2^) indicates decreases in growth rate over time at a temperature. In addition, the interaction between temperature T and t^2^ in the model indicates that the changes in growth rate with time differ among test temperatures; such effects are of particular interest in identifying effects of heat stress or other time-dependent effects.

Second, we modeled mean short-term larval growth rate (g) as a function of test temperature (i.e. the thermal performance curve) for each species, to evaluate how the performance curve changes over time. Here growth rate (g) was defined as the change in mass (Δm) over a time interval (Δt). For *P. rapae* we quantified growth rate of each individual for three time periods: Period 1: 0-24h; Period 2: 24- 48h or 30-54h; and Period 3: 48-80h or 54-80h. For *M. sexta* we also quantified growth rate for three time periods: Period 1: 0-24h; Period 2: 24-48h; and Period 3: 48-72h. We are particularly interested in how the shape of the thermal performance curve for growth rate as a function of test temperature changed with time period.

Our studies with *M. sexta* also determined ingestion (on a dry-mass basis: see above) during each time interval. By summing ingestion across time intervals we obtain cumulative ingestion (C) over time for each individual. Our models for cumulative ingestion and ingestion rate used the same two sets of analyses and time periods described above for body mass. This allows us to directly compare how growth and ingestion rates changed with time across different temperatures, and how performance curves for ingestion rate change with time.

## Results

### A. Growth in *Pieris rapae*

Individual growth curves for *P. rapae* (Fig. 1) show that mass increased almost linearly with time for temperatures between 11 and 23 °C, with more rapid growth (steeper slopes) at 23 °C than at the lower temperatures (Fig. 1). At 23 °C some larvae had completed larval feeding and growth by 54h; at lower temperatures no larvae had completed growth after 80h, and larvae at 11 °C never completed larval growth or pupation. At 29 °C, larval growth was initially rapid but then leveled off after 30-54h. At 35 °C, most larvae initially grew rapidly but then leveled off and declined after 30-54h; some larvae did not grow successfully, and they died within 30h. Most larvae declined in mass at 40-41 °C, and all died within 24h. These patterns illustrate the temporal dynamics of growth change across temperature.

Because we are interested of effects of time and temperature just during the larval growth phase, we restricted our statistical analyses to times 24h or more prior to the initiation of pupation (see Materials and Methods). Mixed-effects models indicated that mass of 5^th^ instar *P. rapae* larvae was significantly affected by time, test temperature and their interaction (Table 1). The mean growth curves at different test temperatures (Fig. 2A) varied both in initial slope (growth rate) and in curvature, with greater (more negative) curvature at temperatures of 29 °C and above. Test temperatures that produced the largest mean size changed with increased exposure time (indicated by the crossing lines in Fig. 2A). As a consequence, the thermal performance curves for growth rate changed with time (Fig. 2B). Mean growth rate was highest at temperatures of 29-35 °C during the 1^st^ time period (0-24h), but by the 2^nd^ time period (24-54h) mean growth rate was highest at 23-29 C (Fig. 2B). By the third time period (48-80h) mean growth rate of the remaining individuals was negative for test temperatures of 29-35 °C. Note that at 29 and 35 °C many larvae had stopped feeding and initiated pupation by 54h; as a consequence, the remaining, slower-growing larvae reduce the estimated mean growth rate for the 3^rd^ time period (see Fig. 1).These results show that for *P. rapae* larvae, growth rates decline over time at the higher temperatures, leading to changes in the thermal performance curve and a reduction in the optimal temperature for growth rate.

**Table 1.** ANOVA results (F-tests) for mass of *Pieris rapae* larvae as a function of time since the start of 5^th^ instar (2^nd^ order polynomial) and test temperature (factor). Initial mass at the start of 5^th^ instar is included as a covariate. The analysis used a linear mixed-effects model with individual as a random effect (intercept), estimating using maximum likelihood. See Methods.

| Term | DF | F | p |
| --- | --- | --- | --- |
| Intercept | 1 | 4073.3 | <0.0001 |
| Time | 2 | 1253.6 | <0.0001 |
| Temperature | 5 | 46.1 | <0.0001 |
| Initial Mass | 1 | 25.0 | <0.0001 |
| Time x Temperature | 10 | 70.8 | <0.0001 |

### B. Growth and ingestion in *Manduca sexta*

Individual growth curves for *M. sexta* (Fig. 3) for test temperatures between 25 and 40 °C show that mass initially increased rapidly, with some acceleration (positive curvature) at temperatures from 25 to 34 °C. At 30 and 34 °C most larvae had begun wandering by 72-96h, whereas at 25 °C few larvae began wandering before 120h (Fig. 3). Growth curves were more strongly (negatively) curved at higher temperatures of 38-40 °C, suggesting declining growth rates over time at these temperatures. At 38 °C wandering did not begin until 96-120h; 87% of larvae reached the wandering phase, but none pupated successfully. At 40 °C most larvae were still alive (but had not wandered) by 144h, but only 53% of larvae reached the wandering phase, and none pupated successfully at this temperature. These results suggest that over short time periods, *M. sexta* larvae can growth rapidly at 38 °C and can maintain positive growth at 40 °C; but longer exposures to these temperatures delay development, slow growth and increase mortality.

Because we are interested of effects of time and temperature just during the larval growth phase, we restricted our statistical analyses to times 24h or more prior to the initiation of wandering (see Materials and Methods). Mixed-effects models indicated that mass of 5^th^ instar *M. sexta* larvae was significantly affected by time, test temperature and their interaction (Table 2A). The mean growth curves at different test temperatures (Fig. 4A) varied both in initial slope (growth rate) and in curvature, with greater negative curvature at higher temperatures during later time periods. As a result, test temperatures that produced the greatest mass gain changed over time (indicated by the crossing lines in Fig. 4A). For example mean mass gain was greatest at 34 °C from 6-54h, but was greatest at 30 °C by 72h. Note that most larvae at 30 and 34 °C had stopped feeding and begun wandering by 96h (Fig. 3), so that mean mass gain for times after 72h are strongly affected by the slower growing larvae that remained (Fig. 3, 4A). As a consequence of these effects, the thermal performance curves for mean growth rate changed with time period (Fig. 4B). Mean growth rate was lower at all test temperatures during the first time period (Fig. 4B) because of small initial mass and the initial acceleration of the growth curves (Fig. 3 and 4A). Mean growth rate was highest at a test temperature of 34 °C during the 1^st^ time period (0-24h) but by the 3^rd^ period (48-72h) mean growth rate was highest at 30 °C (Fig. 2B). These results show that for *M. sexta* larvae, growth rates decline over time at the higher temperatures, leading to changes in the thermal performance curve and a reduction in the optimal temperature for growth rate.

**Table 2.** ANOVA results (F-tests) for mass (A) and cumulative ingestion (B) of *M. sexta* larvae as a function of time since the start of 5^th^ instar (3^rd^ order polynomial) and test temperature (factor). Initial mass at the start of 5^th^ instar is included as a covariate. The analysis used a linear mixed-effects model with individual as a random effect (intercept), estimating using maximum likelihood. See Methods.

| Term | DF | F | p |
| --- | --- | --- | --- |
| Intercept | 1 | 2570.3 | <0.0001 |
| Time | 3 | 1935.3 | <0.0001 |
| Temperature | 4 | 7.9 | <0.0001 |
| Initial Mass | 1 | 34.9 | <0.0001 |
| Time x Temperature | 12 | 44.8 | <0.0001 |

B. Cumulative ingestion
| Term | DF | F | p |
| --- | --- | --- | --- |
| Intercept | 1 | 1253.1 | <0.0001 |
| Time | 3 | 1989.3 | <0.0001 |
| Temperature | 4 | 1.18 | 0.3251 |
| Initial Mass | 1 | 15.0 | <0.0002 |
| Time x Temperature | 12 | 22.0. | <0.0001 |

Individual curves for total (dry-mass) ingestion for *M. sexta* (Fig. 5) were similar to those for growth (Fig. 3) in many respects, but differed in several important ways. Total ingestion accelerated (positive curvature) during the first 2-3 days at test temperatures of 25-30 °C. At all test temperatures from 25- 40 °C, total ingestion continued to increase throughout the measurement period (Fig. 3). In particular at 40 °C, mass of most individuals stopped increasing after ∼ 72h, whereas total ingestion continued to increase until wandering or the end of the measurement period for nearly all individuals.

**Figure 5.**
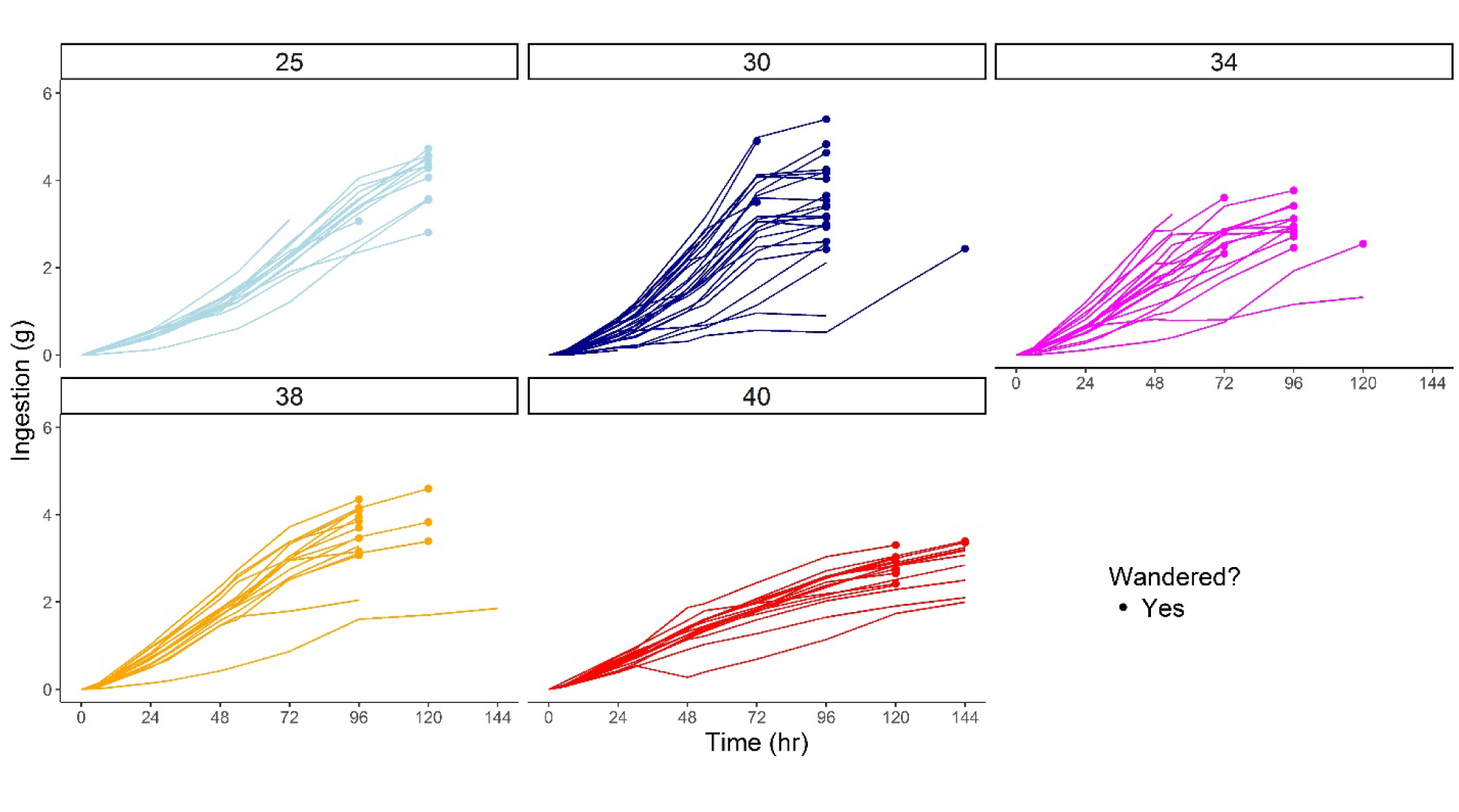
Individual ingestion curves throughout the 5^th^ instar of *M. sexta* by test temperature. Time 0h represents the initial amount eaten (0) upon molting into the final 5^th^ instar and the final measurement represents the total amount eaten during the 5^th^ instar. Individuals who successfully made it through the 5^th^ instar to the wandering stage during the time that measurements took place are denoted with a circle.

Mixed-effects models indicated that total ingestion of 5^th^ instar *M. sexta* larvae was significantly affected by time, test temperature and their interaction (Table 2B). The mean ingestion curves at different test temperatures (Fig. 6A) varied both in slope (growth rate) and curvature. Mean ingestion continued to increase over time even at the highest temperatures, in contrast to mean mass (compare Fig. 6A and 4A). Note that most larvae at 30 and 34 °C had stopped feeding and begun wandering by 96h (Fig. 5), so that mean ingestion for times after 72h are strongly affected by the slower growing (and eating) larvae that remained (Fig. 6A). Ingestion curves had less negative curvature than growth curvatures at all. Thermal performance curves (Fig. 6B) showed that the optimal temperature that maximized ingestion rate decreased from 34-38 °C to 30 °C across the three time periods (Fig. 6B). Compared with growth rate, ingestion rate remained relatively high at the highest temperatures (38-40 °C). These results suggest that time-dependent reductions in performance at high temperatures were greater for growth than for ingestion.

**Figure 6.**
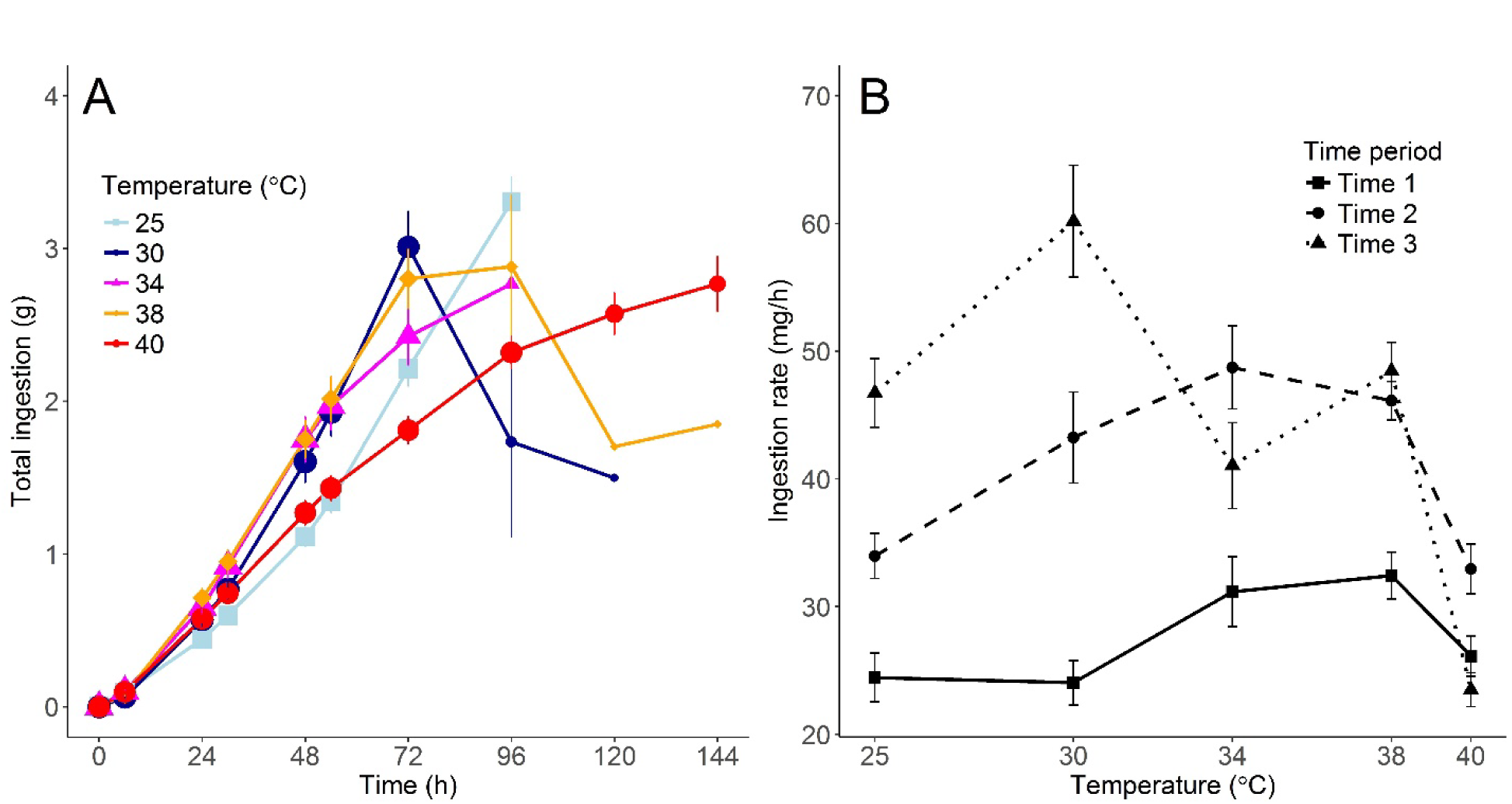
Total mean ingestion (g) with standard error during the 5^th^ instar by time (A) and mean ingestion rate (mg/h) with standard error by test temperature (B) for *M. sexta*. Line types denote 3 different time periods of sampling, including Time 0-24h, 24-48h, or 48-72h of the 5^th^ instar. All measurements prior to wandering are included in the calculation of the means.

## Discussion

The non-linear effects of temperature on rates of growth, ingestion and other biological processes have been abundantly documented (Worner 1992, Lactin et al. 1995, Ruel and Ayres 1999, Tobin et al. 2001, Colinet et al. 2015). Similarly, reductions in performance due to continuing exposure to stressfully high temperatures are widely described (Schulte et al. 2011, Mislan et al. 2014, Rezende et al. 2014). This study examines the relationship between these two phenomena: How continuing thermal exposure changes the thermal sensitivity of growth and ingestion over time.

Our results provide three main findings. First, in agreement with previous studies in these systems (Kingsolver and Woods 1997, Kingsolver et al. 2004), initial larval growth rates during the first 6-24h were greatest at relatively high temperatures, of 29-35 °C for *P. rapae* and ∼ 34 °C for *M. sexta*. This finding supports the general pattern that optimal temperatures for short-term aspects of performance, such as feeding rate, sprint speed or escape rate, are higher than those for longer-term metrics of performance (Huey and Stevenson 1979, Huey and Kingsolver 1989, Angilletta 2009). Second, at high temperatures above the optimum, growth rates decline rapidly over time as a result of stress, but the upper thermal limits differ between the two species. For example, 5^th^-instar *P. rapae* larvae were unable to sustain positive growth and survival to pupation at temperatures above 35 °C. In contrast for *M. sexta*, most 5^th^-instar larvae were able to sustain growth and complete larval development at temperatures of 34-38 °C, and some were successful at temperatures of 40 °C. The greater heat tolerance of *M. sexta* reflects its biogeographic range across Central America and southern North America (Casey 1976, Casey 1977).

Third and most importantly, we document the temporal dynamics of temperature exposure: In particular, how larval growth and ingestion rates decline over time at higher temperatures. For example in *P. rapae*, initial positive growth rates at 29 and 35 °C declines towards (and below) zero growth within 48-80h; a similar pattern of declining growth rates over time occurs at temperatures of 34-40 °C for *M. sexta*. We note that changes in growth rate as defined here (mass gain/time) can occur as a consequence of allometric growth; e.g. the acceleration of growth and ingestion rates observed at lower temperatures in *M. sexta* (Figs. 3 and 5) indicate allometric growth (Sears et al. 2012). However, allometric scaling cannot account for the changes in rank (‘line crossings’) among temperatures across time (Figs. 2A, 4A, 6A): these changes necessarily indicate stress or other time-dependent processes. For *M. sexta*, it is noteworthy that at high temperatures of 38-40 °C, ingestion continues even as growth decline to zero (or below); this suggests that post-ingestive processes are limiting growth at these stressful temperatures (Kingsolver and Woods 1997).

Note that larval growth rates inevitably decline at the end of the growth phase, as larvae approach wandering or pupation (Davidowitz and Nijhout 2004, Nijhout et al. 2006). To focus only on the period of active larval growth, we restricted our statistical analyses of growth and ingestion to times 24h or more before to the initiation of wandering or pupation, and to time periods of 0-54h in *P. rapae* and 0-144h in *M. sexta* (see Materials and Methods). We further restricted our analyses and presentation of thermal performance curves to the first 72-80h of growth (Figs. 2, 4, 6). As a result, our findings are not due to the developmental changes associated with molting and metamorphosis.

There are multiple mechanisms by which continuing exposure to stressful environmental conditions can reduce performance over time (Hofmann 1999, Schulte et al. 2011). For present purposes the key is that these time-dependent effects increase with increasing temperatures, reducing performance (growth and ingestion rates) over time at higher temperatures. As a consequence, the position and shape of the thermal performance curves change over time: in particular the optimal and maximal temperatures decline with duration of exposure (Figs. 2, 4, 6). This is consistent with previous results in these study systems for constant temperatures throughout all of larval development: under these conditions, mean growth rates are maximal at 25-30 °C for *P. rapae* and∼ 30 °C for *M sexta*, and most larvae cannot sustain growth and complete development at 32 °C for *P. rapae* and 35 °C for *M. sexta* (Gilbert 1984, Kingsolver and Nagle 2007). More generally, this supports the pattern that optimal temperatures are lower for long-term than short-term aspects of performance (Angilletta 2009).

The shifts in thermal performance curves documented here occur over a time period of 1-4 days, as a consequence of the cumulative effects of stress. This suggests that a single short (1-4h) exposure to high temperature may have little effect on performance, while repeated diurnal exposure to such temperatures may have substantial negative effects. Previous studies show that *M. sexta* can maintain high rates of larval growth, development and survival when reared in diurnally fluctuating conditions of 25± 10 °C, but performance is substantially reduced when reared at 30±10 °C (Kingsolver et al. 2015): repeated exposure to 40 °C for 2h each day has strong negative consequences. These results have important implications for modeling and predicting the performance consequences of thermal variation and climate change for ectotherms. Multiple studies during the past decade have combined thermal performance curves for fitness (or its components) with climatic data or scenarios to predict the fitness consequences of climate change for insects in different biogeographic regions (Deutsch et al. 2008, Kingsolver et al. 2013, Vasseur et al. 2014). These analyses use performance curves based on fitness estimates at constant temperatures over one or more generations (Frazier et al. 2006), and assume that curves measured under chronic constant conditions can be used to predict mean fitness in fluctuating and changing conditions. Several empirical tests show that this approach that yield poor predictions of mean performance in fluctuating environments, especially at higher mean temperatures (Niehaus et al. 2012, Kingsolver et al. 2015). The current study illustrates non-lethal but stressful conditions can alter the shape and position of thermal performance curves over time scales of 1-4 days, with potentially important consequences for how insects might respond to diurnal fluctuations, heat waves and other patterns of thermal variation. Models that incorporate such time-dependent effects will help in improving our predictions about the performance and fitness consequences of climate change (Kingsolver and Woods 2016).

## Acknowledgements

We thank Kate Augustine, Silvan Goddin, Laura Hamon, Christina Hill, Charlotte Hopson, Katie Massie, Kati Moore, Anna Pearson and Martha Wehling for help with the experiments; James Umbanhowar for suggestions on the statistical analyses; and Art Woods for comments on the manuscript. Research supported in part by NSF awards IBN-0212798 and IOS-152767 to JGK.

## Notes

### Competing Interest Statement

The authors have declared no competing interest.

